# Peri-operative pregabalin does not adversely impact experimental outcomes in rat models of chronic pain

**DOI:** 10.1101/2025.11.19.689268

**Authors:** Francesca Di Domenico, Ryan Patel, Kirsty Bannister

## Abstract

**Introduction:** Multiple mechanisms contribute to the experience of pain where the use of model organisms to dissect mechanistically sensory regulatory circuitry is a vital component of discovering underlying causes of persistent pain in disease states. Such disease states can be modelled in animals using surgical procedures that, ethically, should involve administration of analgesia. However, since basic pain researchers often wish to measure pain-related events, animals may be denied peri-operative analgesia to avoid adversely influencing experimental outcomes.

**Methods:** We conducted a systematic review of peri-operative analgesia usage in rat spinal nerve ligation (SNL) and cancer-induced bone pain (CIBP) models. Using a combination of behavioural testing and in vivo electrophysiology in the dorsal horn of the spinal cord, we assessed the impact of peri-operative pregabalin on nociceptive behaviours in the acute recovery phase, and behavioural and electrophysiological experimental outcomes in the chronic phase, of rat SNL and CIBP models.

**Results:** A literature search revealed that, for studies using rat models of SNL or CIBP, only 5.37 % and 12.69 % respectively reported the use of peri-operative analgesia. We then demonstrated that the use of pregabalin as a peri-operative analgesic reduced mechanical hypersensitivity in the acute period following SNL surgery, with no impact on behavioural, electrophysiological or neuropharmacological outcomes in the chronic phase of either model.

**Conclusions:** This study challenges the basic science researcher’s reasoning that peri-operative analgesia confounds neurobiological outcomes. The use of peri-operative analgesia should be an important consideration to improve animal welfare in chronic models of pain.

**Summary:** Peri-operative analgesia can improve animal welfare without adversely affecting long term behavioural or pharmacological experimental outcomes in chronic pain models.

## 1. Introduction

It is estimated that more than 100 million animals are used in research laboratories around the world each year. Animals involved in scientific procedures frequently experience varying degrees of pain, suffering, and distress, with procedures classified according to severity. In accordance with UK regulations, researchers must implement strategies to minimise pain and discomfort in experimental protocols. Adequate peri-operative care, including the administration of appropriate anaesthetics and analgesics during surgical procedures, is essential in mitigating pain and distress in laboratory animals [15].

In chronic pain models, establishing a persistent pain state is crucial for accurate disease modelling. However, despite the well-documented benefits of peri-operative analgesia, its administration remains inconsistently implemented in the field [7]. Although researchers can often anticipate the type and severity of pain experienced by experimental animals undergoing surgical procedures, concerns about the potential impact of analgesic medications on outcome measures may discourage the implementation of pain-relief regimens [8; 10; 15]. However, identifying the specific reasons for the inconsistent use of peri-operative analgesia is challenging, as they likely vary. Contributing factors may include the absence of regulatory requirements in certain countries and/or reluctance to modify well-established protocols due to concerns about compromising research outcomes. This hesitation is particularly relevant for principal investigators aiming to replicate previous pain models, who may resist refinements out of concern for data comparability.

In clinical practice, peri-operative analgesia is routinely administered to alleviate post-surgical pain. Analgesics used for this purpose are generally classified into two categories: non-opioidergic and opioidergic. Due to the significant adverse effects of opioids, their potential for addiction, and their inability to directly address the underlying cause of pain, alternative peri-operative analgesic strategies have been explored. Over the past decade, pregabalin has gained increasing attention as a viable alternative option [12; 16]. Clinically, peri-operative pregabalin administration has been shown to reduce both pain and opioid consumption during the acute post-operative period, with effects observed up to 48 and 72 hours following thoracotomy [9] and lower limb orthopaedic surgery [5]. While peri-operative analgesia is standard practice in human surgical care, its use in animal models remains inconsistent.

Here, we review the reporting of peri-operative analgesic use in two commonly used rat models of chronic pain, the spinal nerve ligation (SNL) model of neuropathy, and cancer-induced bone pain (CIBP). We additionally investigate whether administering peri-operative pregabalin to SNL and CIBP rats (a) improves behavioural hypersensitivity in the acute post-operative phase and (b) affects experimental outcome measures (behavioural hypersensitivity, neuronal excitability and descending inhibitory signalling) in the chronic phase of the models.

## 2. Methods

### 2.1 Animals

Adult male Sprague-Dawley rats (126 in total) were used for all experiments (Charles River, UK). Animals were group housed (maximum of 5) on a conventional 12:12 h light-dark cycle; food and water were available *ad libitum*. Temperature (20-22 °C) and humidity (55-65 %) of holding rooms were closely regulated. Experimental design/analysis was conducted according to ARRIVE guidelines. All procedures described here were approved by an internal ethics panel and the UK Home Office (licence PP0933098) under the Animals (Scientific Procedures) Act 1986. Only male animals were used to make direct comparisons to reference datasets [1; 11].

### 2.2 Cancer-Induced Bone Pain (CIBP) Model

CIBP was induced as described previously [11]. Briefly, rats weighing 120-140 g were anaesthetised using isoflurane (induction 3 % v/v, maintenance 1.5-2 % v/v) delivered in oxygen (1 l/min). Under aseptic conditions, a small incision was made over the right tibia anterior-medial surface and the bone exposed by blunt dissection. Using a dental drill, a hole was made in the tibia to allow for thin polyethylene tube insertion 1-1.5 cm into the intramedullary cavity. On the day of surgery, flask adherent MRMT-1 cells were released from the flask surface by brief exposure to 0.1 % w/v trypsin-ethylenediaminetetraacetic acid (EDTA) and subsequently pelleted by centrifugation (5 min, 1000 rpm). The pellet was washed with Hanks’ balanced salt solution (HBSS) without calcium, magnesium, or phenol red (Invitrogen, Paisley, UK) and centrifuged again (5 min, 1000 rpm). The MRMT1 cell pellet was suspended in HBSS to a final concentration of 300,000 cells/ml (live cells counted using Tryptan Blue staining, Sigma, Germany) and kept on ice until use. Following each surgery, cell viability was monitored, revealing no more than 10 % cell death following ice-storage (approximately 4 h duration). Using a Hamilton syringe, 3,000 MRMT-1 carcinoma cells in 10 μl HBSS or 10 μl HBSS alone (to generate a sham model) was injected into the cavity. The tube was removed, and the hole plugged with bone restorative material. The surrounding skin was closed with absorbable 4-0 sutures. Rats were given an intraperitoneal injection of either 3 mg/kg pregabalin (PGB; gift from Pfizer) or vehicle (VEH; normal saline). The experimenter was blinded to the injection material (MRMT-1 cells or HBSS) during surgery and peri-operative analgesia.

### 2.3 Spinal Nerve Ligation (SNL) Model

SNL was performed as described previously [1]. Briefly, rats weighing 120-140 g were anaesthetised using isoflurane (induction 3 % v/v, maintenance 1.5-2 % v/v) delivered in oxygen (1 l/min). Under aseptic conditions a paraspinal incision was made and the tail muscle retracted from the spinal column. Part of the L5 transverse process was removed to expose the left L5 and L6 spinal nerves, which were then isolated with a glass nerve hook and ligated with a non-absorbable 6-0 braided silk thread proximal to the formation of the sciatic nerve. The surrounding skin and muscle was closed with absorbable 4-0 sutures. Sham surgery was performed in an identical manner omitting the nerve isolation and ligation step. Rats were given an intraperitoneal injection of either 3 mg/kg pregabalin (PGB; gift from Pfizer) or vehicle (VEH; normal saline).

### 2.4 Behavioural tests

All behavioural assessments were conducted by a single experimenter at a consistent time of day during the light phase. Animals were allowed to habituate to handling, the experimenter and apparatus before all behavioural testing. For assessment of mechanical withdrawal thresholds, following room acclimatisation (1 h), rats were placed in isolation inside Perspex chambers on a wire mesh floor and left to acclimatise for a further 15 min. Mechanical sensitivity was assessed using von Frey filaments (Touch-Test, North Coast Medical, USA). Filaments were applied to the plantar hind paw of the rat until they buckled for 5-6 s. Lifting, flinching, and shaking were considered positive responses. Fifty per cent withdrawal thresholds were determined using the up-down method described previously [4], using von Frey filaments with bending forces of 1.4 g, 2 g, 4 g, 6 g, 8 g, 10 g and 15 g. Testing began at 4 g followed by the next weight up or down depending on a negative or positive response respectively. Following a change in direction of the response a further 4 filaments, with a 2 min inter-stimulus recovery period between each stimulation. Paw withdrawal thresholds (50% PWT) were calculated with the following formula: 50% PWT = (10^(x + kδ)/10,000), where x represents the log of the last von Frey tested, δ represents the mean difference between the von Frey filaments in log units (0.17) and k, a value dependent on the series of responses.

Weight bearing was assessed using an incapacitance tester (Linton Instrumentation, Norfolk, UK) in which rats were placed in a plexiglass enclosure where each hind paw resides on a separate weighing plate. Once the animal acquired a relaxed position, the mass exerted by each hind paw was measured 5 times (expressed in grams), with a resting period between measurements to allow the animal to re-equilibrate or slightly shift weight distribution (around 10-20 s). Measurements from each paw were averaged separately and results were transformed to give the percentage of weight borne on each side to the total rear leg bearing (taken as 100%).

### 2.5 In Vivo Electrophysiology

*In vivo* electrophysiology was performed days 14-15 post-CIBP induction and days 10-14 post-SNL induction (in 250-300 g rats). Anaesthesia was initially induced with 3.5% v/v isoflurane delivered in 3:2 ratio of nitrous oxide and oxygen. Once areflexic, a tracheotomy/canulation was performed and rats were subsequently maintained on 1.5% v/v isoflurane for the remainder of the experiment (approximately 3-4 h; core body temperature was maintained throughout with the use of a homeothermic blanket). Rats were secured in a stereotaxic frame and a laminectomy was performed to expose the L4-L6 segments of the spinal cord; two spinal clamps were applied to stabilise the spinal column. Extracellular recordings were obtained from deep dorsal horn wide dynamic range lamina V/VI neurones with receptive fields on the glabrous skin of the hind toes using 127 µm diameter 2 MΩ parylene-coated tungsten electrodes (A-M Systems, Sequim, WA). The search stimulus consisted of light tapping of the hind paw as the electrode was manually lowered. Neurones were characterised from depths relating to the deep dorsal horn laminae, and once a single unit was isolated neurones were classified as wide dynamic range on the basis of sensitivity to dynamic brushing, noxious mechanical (60 g) and noxious heat stimulation (48 °C) of the receptive field. Data were captured and analysed by a CED Micro1401 interface coupled to a computer with Spike2 v4 software (Cambridge Electronic Design, Cambridge, United Kingdom). The signal was amplified (pre-amp+amp x30-40k), bandpass filtered (low/high frequency cut-off 1/3 kHz) and digitised at rate of 20 kHz. A HumBug (Quest Scientific, Canada) was used to remove low frequency noise (50-60 Hz).

Each baseline trial involved applying von Frey filaments (8 g, 26 g and 60 g) to the receptive field for 10 s per stimulus (60 s recovery period between stimuli). Evoked responses were quantified as the total number of neuronal events during the 10 s stimulation. Once stable neuronal responses were obtained (three consistent recordings <10 % variation in response), a diffuse noxious inhibitory controls (DNIC) test was applied. DNIC were activated by a noxious clamp (a 35 mm bulldog serrefine (Interfocus, Linton, UK)) applied to the ipsilateral ear concurrently to stimulation of the hind paw with von Frey filaments. A 10 min non-stimulation recovery period was allowed before two more baseline trials were conducted (baseline responses were calculated as a mean of three baseline trials). Atipamezole (100 μg/50 μl; vehicle: 97 % normal saline/2 % cremophor/1 % DMSO. Sigma-Aldrich, Gillingham, UK) was subsequently applied topically to the spinal cord and tests repeated at 10, 20, 30, 60 min post-dosing; peak change from baseline is plotted.

### 2.6 Statistics

Statistical analyses were performed using SPSS v29 (IBM, Armonk, NY). The experimental unit for behaviour and *in vivo* electrophysiology was the individual rat. Rats were pseudorandomised into experimental/treatment groups by an independent investigator using a random number generator. All behavioural experiments were conducted blind to experimental variables; unblinding was performed after data collection. Group sizes were determined by *a priori* calculations using the following assumptions: F test, α 0.05, 1-β 0.8, ε 1, effect size range *d*=0.5 to 0.8 for electrophysiological measures, effect size range *d*=0.3 to 0.5 for behavioural measures. Mechanical coding of neurones were compared with a 2-way repeated measures (RM) ANOVA, followed by a Bonferroni *post hoc* test for paired comparisons. Where appropriate, sphericity was tested using Mauchly’s test; the Greenhouse-Geisser correction was applied if violated. Main effect of DNIC or drug is reported throughout. Mechanical hypersensitivity and weight bearing were assessed with either a Friedman test or Mann-Whitney. Effect sizes are reported as Cohen’s *d*. All data plotted represent mean ± SEM, \**P*<0.05, *P*<0.01,\*\*\**P*<0.001.

## 3. Results

### 3.1 A review of the literature: the frequency of peri-operative analgesic use in SNL and CIBP models

The study selection details were based on a PubMed database, the search identified 1183 papers from 1976 until 2022 that cited the use of animals undergoing CIBP or SNL surgery (Fig. 1).

**Figure 1:**
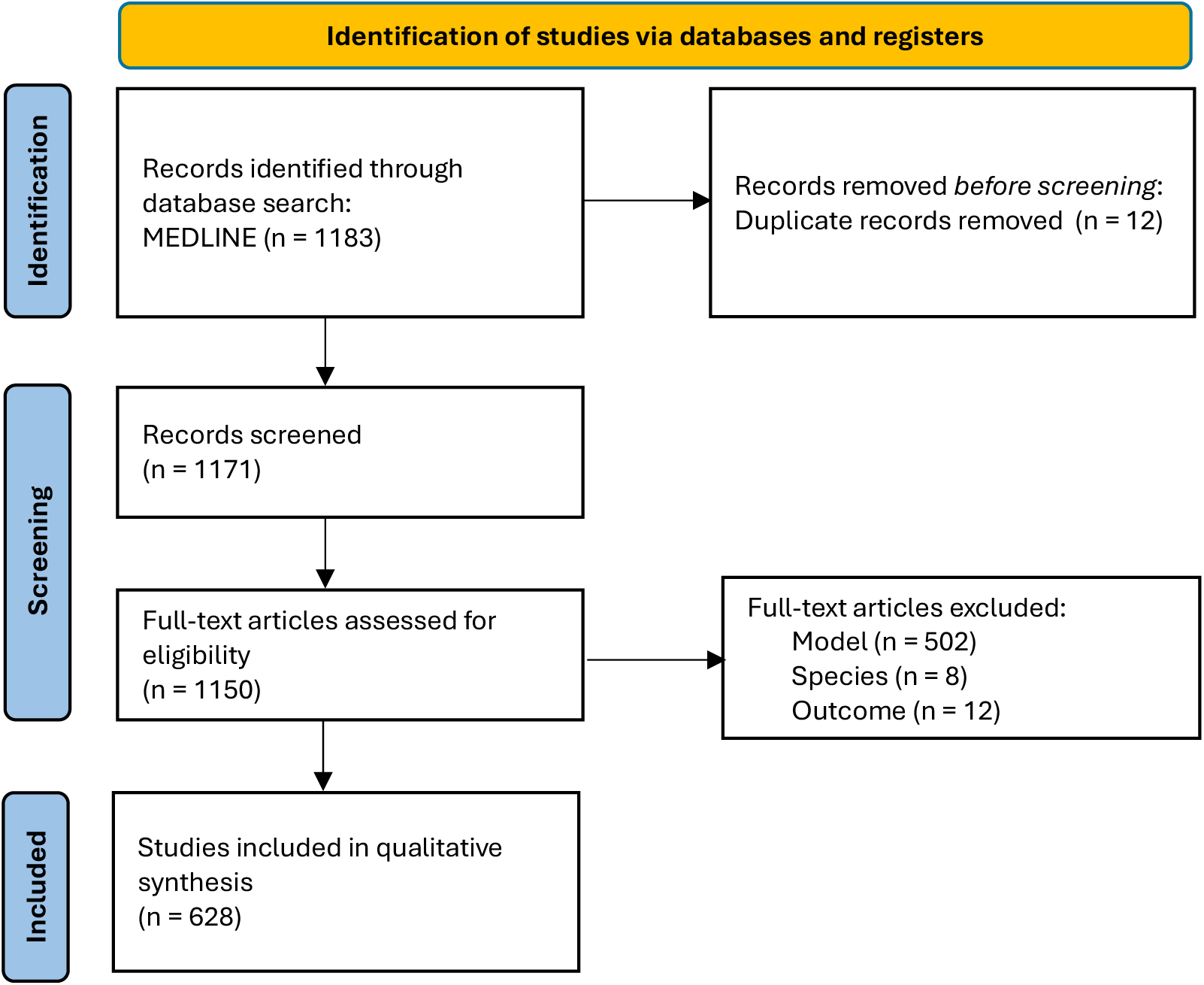
PRISMA Flow Diagram. This diagram shows the systematic process we followed to include papers captured by our search. Search terms (Species: rat, murine. Intervention: analgesic, spinal nerve ligation, cancer induced bone pain. Outcome: behavior, in vivo electrophysiology, calcium imaging). Adapted from [13] under a creative commons licence CC BY 4.0.

For the SNL model we found 1012 papers that cited the use of spinal nerve ligation of which 498 papers were excluded from the study (Table 1). Exclusion criteria included species (only rats were included), neuropathic pain model (only rodents that had undergone spinal nerve ligation were included) and experimental outcome (only studies that included a behavioural measure were included). Thus, information was collated from 502 studies. The analysis revealed that of the 502 original research articles, only 5.37% (27 studies) reported the use of peri-operative analgesia. The remaining 475 did not report the use of peri-operative analgesia.

**Table 1:** Details from papers that reported the use of peri-operative analgesia in SNL, with particular reference to the experimental approach, anaesthesia and analgesia used.

|  |  |
| --- | --- |
| SNL PAPERS INCULDED (502) |  |
| PERI-OPERATIVE ANALGESIA (27) | ANALGESIA |
|  | MELOXICAM (1) |
|  | KETAMINE (21) |
|  | BUPRENORPHINE (1) |
|  | ZOLETIL (2) |
|  | MIDAZOLAM/HYPNORM (1) |
|  | MEDETOMIDINE (2) |

We found 171 papers that cited the use of a rodent model of CIBP, of which 45 papers were excluded from the study (Table 2). Exclusion criteria included species (only rats were included), cancer model (only rodent models of cancer induced bone pain were included) and experimental outcome (only studies that included a behavioural measure were included). Thus, information was collated from 126 studies. The analysis revealed that of the 126 original research articles, only 12.69% (16 studies) reported the use of peri-operative analgesia. The remaining 113 articles did not report the use of peri-operative analgesia.

**Table 2:**
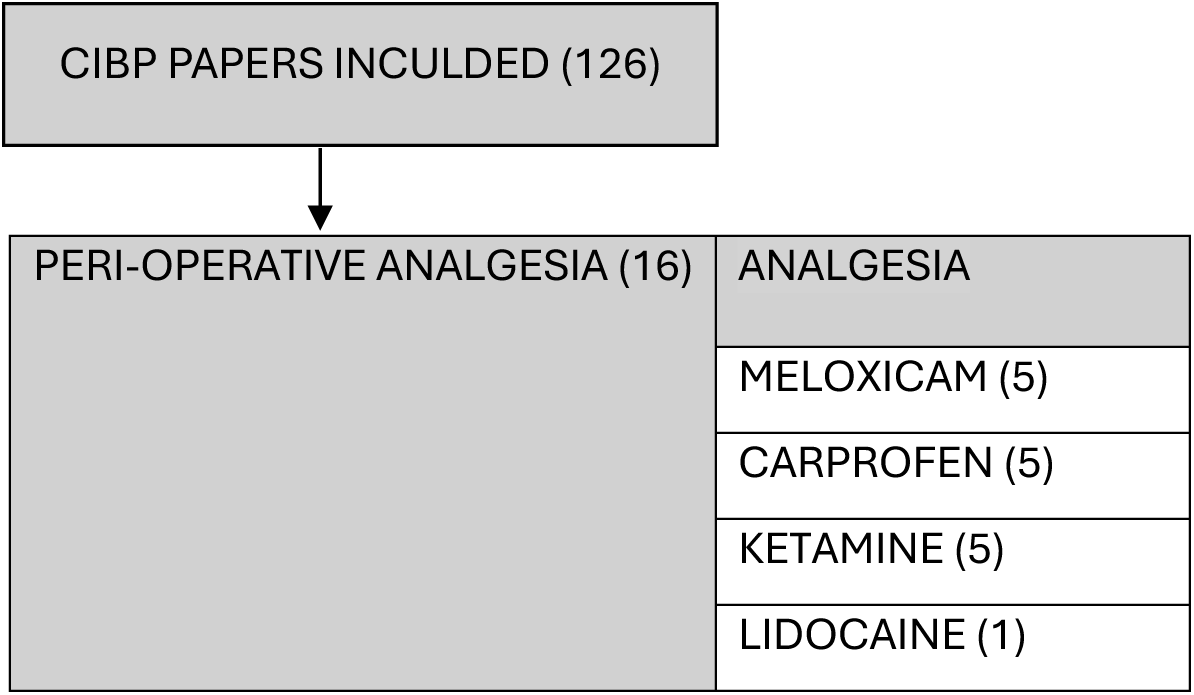
Details from papers that reported the use of peri-operative analgesia in CIBP, with particular reference to the experimental approach, anaesthesia and analgesia used.

|  |  |
| --- | --- |
| CIBP PAPERS INCULDED (126) |  |
| PERI-OPERATIVE ANALGESIA (16) | ANALGESIA |
|  | MELOXICAM (5) |
|  | CARPROFEN (5) |
|  | KETAMINE (5) |
|  | LIDOCAINE (1) |

### 3.2 SNL, but not CIBP rats, showed a mild improvement in mechanical hypersensitivity in the acute post-surgical phase when pregabalin was administered as a peri-operative analgesic

We examined the effect of peri-operative pregabalin on the development of primary (SNL) and secondary (CIBP) mechanical hypersensitivity during the acute post-operative phase defined as days 1-7 post-injury; sham animals (sham^s^ and sham^c^ respectively) were included as control groups which experienced acute tissue damage but no chronic injury. There was no change from baseline of mechanical withdrawal thresholds in the sham^s^ group given peri-operative vehicle, but we observed a decrease in the pregabalin group (Friedman test: sham^s^ (VEH) *P*=0.135, sham^s^ (PGB) *P*=0.00003) (Fig. 2A). Both SNL groups rapidly developed mechanical hypersensitivity from day 1 onwards (Friedman test: SNL (VEH) *P*=0.00002, SNL (PGB) *P*=0.000005) (Fig. 2B), with evidence of higher withdrawal thresholds in SNL rats receiving peri-operative analgesia on day 3 post-injury (Mann-Whitney, day 2: *P*=0.091, day 3: *P*=0.00054, day 7: *P*=0.088). Sham^s^ rats treated with pregabalin did not exhibit any time dependent changes in weight bearing with weak evidence for altered weight bearing observed in vehicle treated rats (Friedman test: sham^s^ (VEH) *P*=0.022, sham^s^ (PGB) *P*=0.52) (Fig. 2C). Both SNL groups avoided placing weight on the injured paw from day 1 onwards (Friedman test: SNL (VEH) *P*=0.00006, SNL (PGB) *P*=0.00003) (Fig. 2D), with evidence of greater avoidance in SNL rats not receiving peri-operative analgesia on day 2 post-injury (Mann-Whitney, day 2: *P*=0.021, day 3: *P*=0.15).

**Figure 2.**
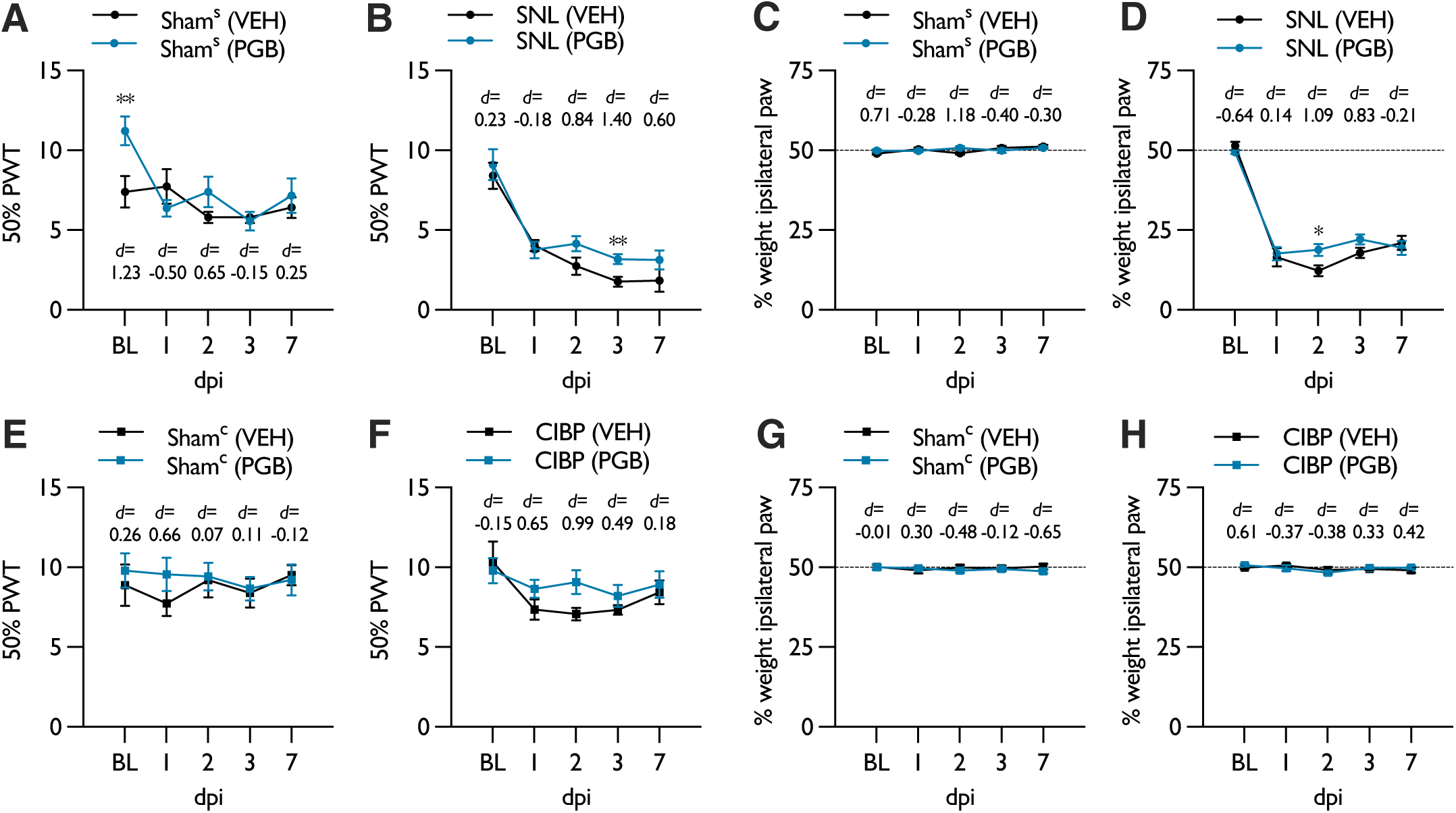
Post-surgical outcomes: effect of peri-operative pregabalin on mechanical hypersensitivity in SNL and CIBP rats during the acute post-operative phase. Development of mechanical hypersensitivity in **(A)** sham^s^ (*n_VEH_*=10, *n_PGB_*=12) and **(B)** SNL rats (*n_VEH_*=10, *n_PGB_*=12). Changes in weight bearing in **(C)** sham^s^ and **(D)** SNL rats. Development of mechanical hypersensitivity in **(E)** sham^c^ (*n_VEH_*=9, *n_PGB_*=9) and **(F)** CIBP rats (*n_VEH_*=10, *n_PGB_*=12). Changes in weight bearing in **(G)** sham^c^ and **(H)** CIBP rats. Data represent mean ± SEM; *P<0.05, **P<0.01. Asterisks denote dicerence between vehicle and pregabalin treated groups. (BL – baseline, CIBP – cancer-induced bone pain, dpi – days post injury, PGB – pregabalin, SNL – spinal nerve ligation, VEH - vehicle).

There was no change from baseline of mechanical withdrawal thresholds in sham^c^ groups given peri-operative vehicle or pregabalin (Friedman test: sham^c^ (VEH) *P*=0.43, sham^c^ (PGB) *P*=0.93) (Fig. 2E). Neither CIBP groups developed secondary mechanical hypersensitivity (Friedman test: CIBP (VEH) *P*=0.12, CIBP (PGB) *P*=0.42) (Fig. 2F). Weight bearing was not altered in sham^c^ (Friedman test: sham^c^ (VEH) *P*=0.32, sham^c^ (PGB) *P*=0.45) (Fig. 2G) or CIBP rats (Friedman test: CIBP (VEH) *P*=0.25, CIBP (PGB) *P*=0.21) (Fig. 2H). Heat hypersensitivity was not observed in SNL or CIBP rats (data not shown).

### 3.3 Administration of peri-operative pregabalin does not adversely impact nocifensive behaviours in the chronic phase of SNL and CIBP models

We examined whether administering peri-operative analgesia affected experimental outcomes in the chronic phase of the models. At day 14 post-injury, SNL rats developed primary mechanical hypersensitivity (Mann-Whitney, VEH: *P*=0.00003, PGB: *P*=0.000005) (Fig. 3A) and exhibited altered weight bearing (Mann-Whitney, VEH: *P*=0.00001, PGB: *P*=0.000001) (Fig. 3B), however no difference in the magnitude of primary mechanical hypersensitivity or weight bearing was observed between rats receiving peri-operative vehicle or pregabalin. Similarly, CIBP rats developed secondary mechanical hypersensitivity (Mann-Whitney, VEH: *P*=0.0005, PGB: *P*=0.000007) (Fig. 3C) and exhibited altered weight bearing (Mann-Whitney, VEH: *P*=0.00008, PGB: *P*=0.00006) (Fig. 3D), however no difference in the magnitude of secondary mechanical hypersensitivity or weight bearing was observed between rats receiving peri-operative vehicle or pregabalin.

**Figure 3.**
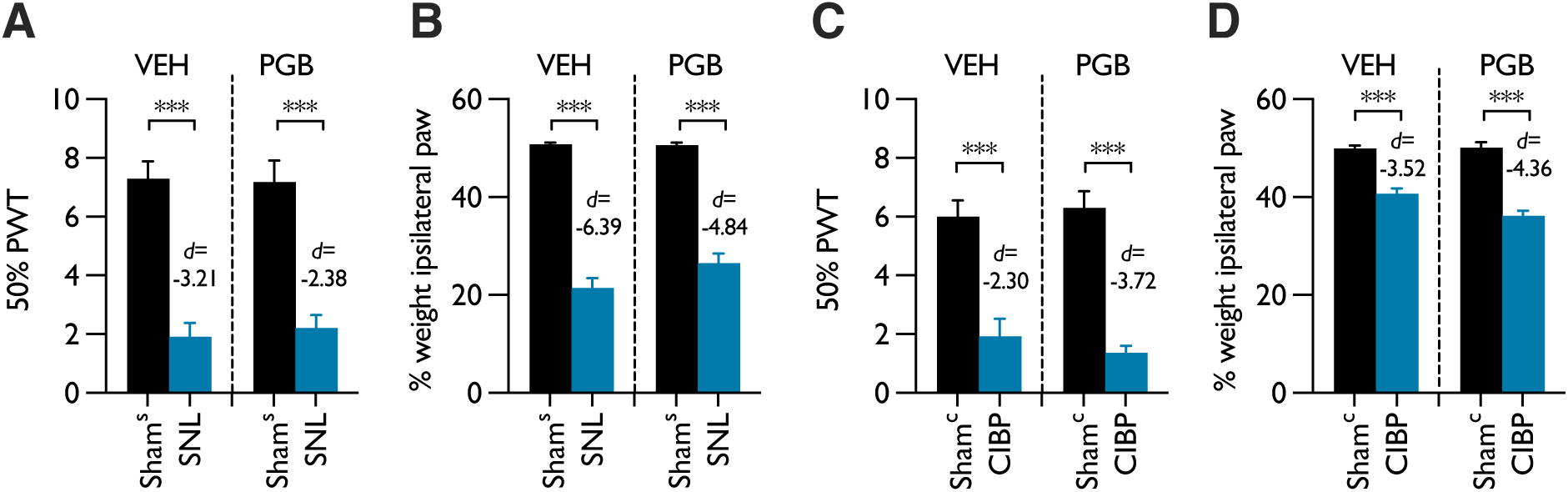
Experimental outcomes (*i*): effect of peri-operative pregabalin on the establishment of chronic pain in SNL and CIBP rats. **(A**) Effect of peri-operative pregabalin on mechanical hypersensitivity in sham^s^ (*n_VEH_*=10, *n_PGB_*=12) and SNL rats (*n_VEH_*=10, *n_PGB_*=12) on day 14 post-injury. **(B)** Effect of peri-operative pregabalin on weight bearing in sham^s^ and SNL rats on day 14 post-injury. **(C)** Effect of peri-operative pregabalin on mechanical hypersensitivity in sham^c^ (*n_VEH_*=9, *n_PGB_*=9) and CIBP rats (*n_VEH_*=10, *n_PGB_*=12) on day 14 post-injury. **(D)** Effect of peri-operative pregabalin on weight bearing in sham^c^ and CIBP rats on day 14 post-injury. Data represent mean ± SEM; \*\*\**P*<0.001. (CIBP – cancer-induced bone pain, PGB – pregabalin, SNL – spinal nerve ligation, VEH - vehicle).

### 3.4 Administration of peri-operative pregabalin does not impact the expression of descending inhibition in SNL and CIBP rats

We examined whether administering peri-operative analgesia affected endogenous pain modulation in the chronic phase of the models. *In vivo* electrophysiology was performed in the dorsal horn of the spinal cord to record from wide dynamic range neurones in lamina V/VI. First, descending inhibition was assessed by activating diffuse noxious inhibitory controls (DNIC) which were recruited by applying a distant noxious stimulus during hind paw stimulation. DNIC were expressed in sham^s^ rats in peri-operative vehicle (2-Way RM ANOVA: *P*=0.02, *F*_1,4_=50.952) and pregabalin treated groups (2-Way RM ANOVA: *P*=0.000007, *F*_1,4_=286.94) (Fig. 4A). DNIC were absent in both peri-operative vehicle and pregabalin treated SNL rats (2-Way RM ANOVA: VEH - *P*=0.804, *F*_1,4_=0.07; PGB - *P*=0.451, *F*_1,7_=0.636) (Fig. 4B). DNIC were expressed in sham^c^ rats in peri-operative vehicle (2-Way RM ANOVA: *P*=0.001, *F*_1,4_=74.752) and pregabalin treated groups (2-Way RM ANOVA: *P*=0.0034, *F*_1,3_=72.077) (Fig. 4C). DNIC were expressed in CIBP rats in peri-operative vehicle (2-Way RM ANOVA: *P*=0.0002, *F*_1,4_=172.075) and pregabalin treated groups (2-Way RM ANOVA: *P*=0.0032, *F*_1,4_=39.95) (Fig. 4D).

**Figure 4.**
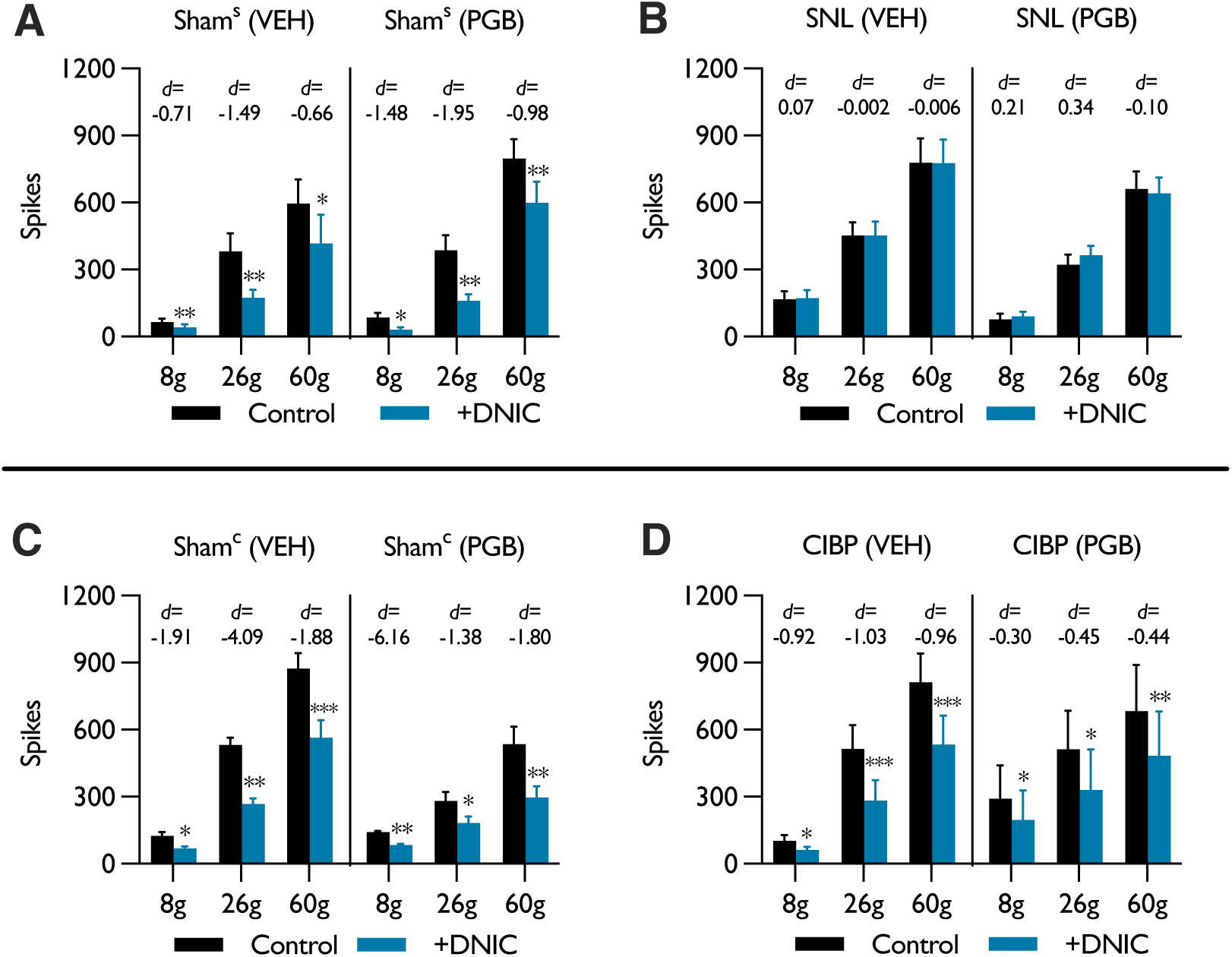
Experimental outcomes (*ii*): effect of peri-operative pregabalin on the expression DNIC in SNL and CIBP rats. Effect of activating DNIC on neuronal responses to mechanical stimulation in **(A)** sham^s^ (*n_VEH_*=5, *n_PGB_*=5) and **(B)** SNL rats (*n_VEH_*=5, *n_PGB_*=8). Effect of activating DNIC on neuronal responses to mechanical stimulation in **(C)** sham^c^ (*n_VEH_*=5, *n_PGB_*=4) and **(D)** CIBP rats (*n_VEH_*=5, *n_PGB_*=5). Data represent mean ± SEM; \**P*<0.05, \*\**P*<0.01, \*\*\**P*<0.001. (CIBP – cancer-induced bone pain, DNIC – diffuse noxious inhibitory controls, PGB – pregabalin, SNL – spinal nerve ligation, VEH - vehicle).

Subsequently, noradrenergic inhibitory tone was revealed by applying the α_2_-adrenoceptor antagonist atipamezole topically to the spinal cord. Neuronal responses to mechanical stimulation were facilitated in sham^s^ rats in peri-operative vehicle (2-Way RM ANOVA: *P*=0.022, *F*_1,4_=13.168) and pregabalin treated groups (2-Way RM ANOVA: *P*=0.0495, *F*_1,4_=7.762) (Fig. 5A). Compared to sham^s^ rats, inhibitory tone was reduced in SNL rats in both the peri-operative vehicle (2-Way RM ANOVA: *P*=0.001, *F*_1,4_=70.259) and pregabalin treated groups (2-Way RM ANOVA: *P*=0.730, *F*_1,4_=0.137) (Fig. 5B). Neuronal responses were unaffected in sham^c^ rats post-atipamezole dosing in groups treated with peri-operative vehicle (2-Way RM ANOVA: *P*=0.751, *F*_1,4_=0.115) and pregabalin (2-Way RM ANOVA: *P*=0.08, *F*_1,3_=6.775) (Fig. 5C). Neuronal responses in CIBP rats were unaffected post-dosing of atipamezole in the peri-operative vehicle treated group (2-Way RM ANOVA: *P*=0.815, *F*_1,4_=0.63), but weak evidence was found for altered neuronal responses in the pregabalin treated group (2-Way RM ANOVA: *P*=0.005, *F*_1,4_=32.672, paired comparisons *P*>0.05) (Fig. 5D). Effect sizes were broadly comparable across all groups.

**Figure 5.**
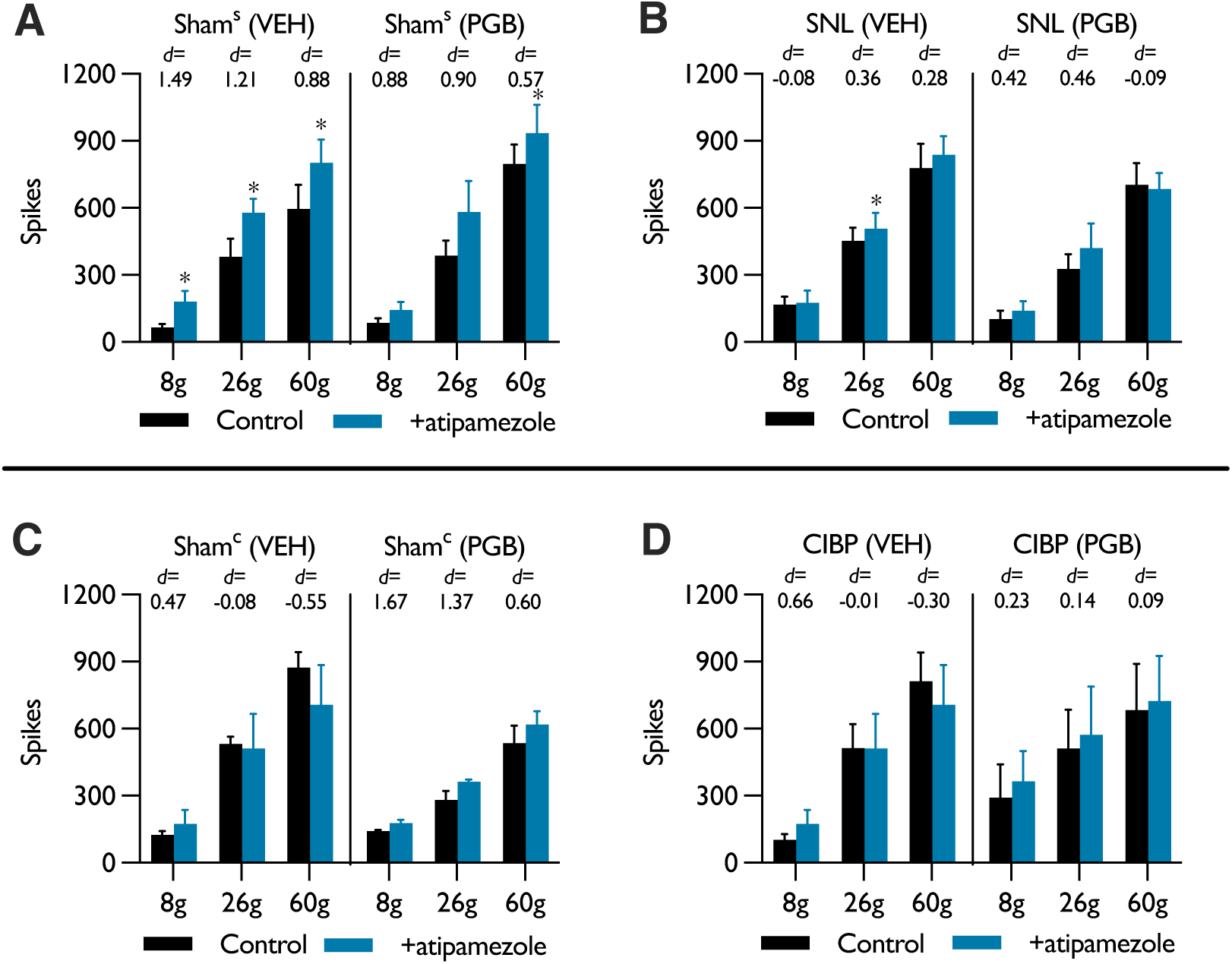
Experimental outcomes (*iii*): effect of peri-operative pregabalin on noradrenergic inhibitory tone in SNL and CIBP rats. Effect of spinal atipamezole on neuronal responses to mechanical stimulation in **(A)** sham^s^ (*n_VEH_*=5, *n_PGB_*=5) and **(B)** SNL rats (*n_VEH_*=5, *n_PGB_*=5). Effect of spinal atipamezole on neuronal responses to mechanical stimulation in **(C)** sham^c^ (*n_VEH_*=5, *n_PGB_*=4) and **(D)** CIBP rats (*n_VEH_*=5, *n_PGB_*=5).Data represent mean ± SEM; \*\*\**P*<0.001. (CIBP – cancer-induced bone pain, PGB – pregabalin, SNL – spinal nerve ligation, VEH - vehicle).

## 4. Discussion

A significant proportion of published original research articles that include the use of rats undergoing surgery for spinal nerve ligation or cancer induced bone pain models do not cite the use/administration of peri-operative analgesia. This was unexpected, not least since, in the UK, the use of analgesics during surgery is stipulated, for procedures that are classified as moderate to severe, as a necessity by the regulatory bodies. It is noteworthy that the umbrella of dates captured by our search began as early as 1976. One would expect that, proportionally, the number of studies citing the use of peri-operative analgesics will increase as time goes on, particularly in countries where animal welfare is governed and/or informed by bodies such as the NC3Rs. Nonetheless, our findings demonstrate that historically as well as recently, animal welfare has not always been reported in research papers for a significant number of animals undergoing surgeries to induce a chronic pain state.

In our study pregabalin was chosen as the peri-operative analgesic for several reasons. First, due to its use clinically as an opioid replacement during surgical procedures [6], second due to its known peripheral and spinal mechanism of action in neuropathic pain [3; 14], and third due to its modulation of affective-motivational qualities of pain [2]. A key consideration for interpreting the behavioural data is the application of tests to the hind paw in both models. For the SNL model this is the injured territory of the nerve and represents primary mechanical hypersensitivity, whereas for the CIBP model this represents secondary mechanical hypersensitivity. We did not observe secondary hypersensitivity in the CIBP model during the acute post-operative phase, and pregabalin did not affect the development of secondary mechanical hypersensitivity at day 14 of the model. In contrast, primary mechanical hypersensitivity was evident at day 1 post-injury in the SNL model and peri-operative pregabalin provided mild relief on days 2 and 3 post-injury without affecting behavioural responses in the chronic phase of the model. Collectively these observations suggest peri-operative pregabalin improves primary pain after injury without affecting the establishment of primary or secondary hypersensitivity in the chronic phase. With a single dose, the precise mechanism of action of peri-operative pregabalin is unclear. Chronic dosing is known to reduce peripheral axonal trafficking of α2δ-1 subunits [3] but this mechanism cannot account for the effects observed here which are suggestive of an acute/transient central mechanism of action.

After demonstrating that behavioural outcomes were not affected by peri-operative analgesia, we replicated two previous electrophysiological studies of descending inhibition in the SNL and CIBP models. These previous studies did not administer peri-operative analgesics allowing us to draw comparisons with these reference datasets. Firstly, we show here that the expression of an endogenous modulatory circuit, diffuse noxious inhibitory controls (DNIC), was not impacted by peri-operative pregabalin. We have previously observed an absence of DNIC in the SNL model [1] and preservation of DNIC in the CIBP model [11] at comparable timepoints. Secondly, we investigated tonic noradrenergic inhibitory tone and observed that the effects of spinal atipamezole on neuronal excitability was consistent between peri-operative vehicle and pregabalin groups, and in line with our previous observations [1; 11]. These data indicate that peri-operative pregabalin does not adversely affect this particular pathophysiological mechanism and the functionality of endogenous pain modulatory pathways in the chronic phase of SNL and CIBP models.

In summary, our data support that peri-operative analgesia can improve animal welfare in the acute post-operative phase without affecting desirable outcomes in the chronic phase. Developing a standardised framework for peri-operative analgesia is likely to be challenging and must be adapted according to the experimental objectives. For example, researchers investigating the development of a chronic pain state would be negatively impacted by peri-operative analgesia. However, most studies are based on an established chronic pain state as this reflects the clinical situation. Peri-operative analgesic regimens for chronic pain models should be explored more widely and factored into experimental design as refinement of models is possible without detriment to studying pathophysiological mechanisms, contrary to historical dogma.

## Author contributions

All authors contributed to data analysis/interpretation and drafting/editing of the manuscript. FDD conducted experiments, and FDD and KB contributed to study design.

## Abbreviations

CIBP: cancer-induced bone pain
BL: baseline
DNIC: diffuse noxious inhibitory controls
dpi: days post-injury
PGB: pregabalin
PWT: paw withdrawal threshold
RM: repeated measures
SNL: spinal nerve ligation
VEH: vehicle
WDR: wide dynamic range

